# The production of within-family inequality: Insights and implications of integrating genetic data

**DOI:** 10.1101/2020.06.06.137778

**Authors:** Jason M. Fletcher, Yuchang Wu, Zijie Zhao, Qiongshi Lu

## Abstract

The integration of genetic data within large-scale social and health surveys provides new opportunities to test long standing theories of parental investments in children and within-family inequality. Genetic predictors, called polygenic scores, allow novel assessments of young children’s abilities that are uncontaminated by parental investments, and family-based samples allow indirect tests of whether children’s abilities are reinforced or compensated. We use over 16,000 sibling pairs from the UK Biobank to test whether the relative ranking of siblings’ polygenic scores for educational attainment is consequential for actual attainments. We find strong evidence of compensatory processes, on average, where the association between genotype and phenotype of educational attainment is reduced by over 20% for the higher-ranked sibling compared to the lower-ranked sibling. These effects are most pronounced in high socioeconomic status areas. We find no evidence that similar processes hold in the case of height or for relatives who are not full biological siblings (e.g. cousins). Our results provide a new use of polygenic scores to understand processes that generate within-family inequalities and also suggest important caveats to causal interpretations the effects of polygenic scores using siblingdifference designs.

## Introduction

A large social science literature has produced theoretical and empirical support that parental actions shape patterns of within-family inequalities over the life cycle (1–3). Theoretically, the key components of the question include parental attitudes about inequalities of outcomes of their children (4, 5) and the potential of differential returns to investment based on children’s talents (i.e. technologies of skill formation)(6–8). Separating these mechanisms has proven difficult. Empirically, measurement limitations have been important bottlenecks in progress. Ideally, researchers could use measures that (a) occur early in life so they represent endowments and (b) capture endowments of children that are not related to parental behaviors and investments (no feedback effects). Typically, birth weight has been used in analyses that examine whether parents reinforce or compensate for children’s endowments (3, 9–11). However, focusing on birth weight is imperfect because it can be affected by parental behaviors and investments (12–16) and it limits the scope of analysis due to a focus on a single measurement. This scope limitation occurs both in terms of life course outcomes that can be tied to birth weight as well as failing to examine parental responses to endowments that are not associated with birthweight. Alternative measures, such as test scores (4) can be problematic because parents can shape these outcomes prior to research measurement and they may not be “early enough” to capture endowments—for example, before children acquire language skills.

Summary genetic assessments (i.e. polygenic scores, PGS) have the capacity to overcome these empirical limitations, as these measures are fixed at conception—thus, they have no feedback effects and they can capture endowments tied to early outcomes. PGS also provide the possibility of expanding the domains of analysis outside of birthweight. Studies have begun to use these measures to show associations with early childhood outcomes (17), and some research has found evidence that PGS are associated with parental responses (18, 19). However, fewer studies have incorporated these measures into analyses of within-family inequalities. Might parents use observable phenotypic downstream outcomes tied to PGS in their efforts to increase or decrease differences in later outcomes of their children? We begin this direction of empirical analysis by linking two highly predictive PGS measures (education and height) to phenotypic outcomes in adulthood in a sample of over 16,000 sibling pairs from the UK Biobank.

We propose that theoretical models of parental responses to children’s endowments can be assessed with PGS measures of siblings. Because our data do not include measures of early life, we use an indirect test of the accumulated parental responses to their children’s abilities by comparing educational attainments of the siblings with their PGS measurements. In order to provide an omnibus test of compensation vs. reinforcement, we test whether the *relative ranking* of PGS within sibling pairs is consequential for predicting adult outcomes. We focus on the case of education as an exemplar where parents may have the means and desire to shape inequalities in their children’s attainments (20) and the case of height, where they have neither, so that we have a negative test. We hypothesize that if parents prefer, on average, for equalizing their children’s outcomes (i.e. compensatory behaviors), the PGS will be less predictive of attainment in the sibling with a higher relative rank. If parents prefer, on average, to reinforce their children’s relative advantages, the PGS will be more predictive of attainment in the sibling with a higher relative rank. We then explore whether these patterns differ by the socioeconomic status of the families. A large literature has shown that parents in advantaged settings often show compensatory preferences and behaviors while parents in disadvantaged settings show reinforcing preferences and behaviors (11, 21, 22). Therefore, we divide the families in our data based on area-level socioeconomic status and estimate the relative ranking effects outlined above.

## Results

We analyzed educational attainment in the full sample and a sibling sub-sample from respondents with European ancestry in the UK Biobank. Siblings were matched based on kinship estimates of genetic relatedness (see Methods). SI Appendix Table A1 provides descriptive statistics of the full sample and our sibling analysis sample. Our baseline results conformed with other analyses (23–25) showing that the education attainment (EA)-PGS predicts educational attainment in the UKB sample (Table 1, Column 1), that these associations are retained in the sibling sub-sample (Table 1, Column 2) and that the association is reduced by >50% when family fixed effects are included (Table 1, Column 3). Specifically, the association between a standard deviation increase in EA-PGS and EA falls from 0.91 years in Column 1 to 0.44 years in Column 3.

**Table 1.** Associations between PGS-Education and Educational Attainment Comparing Between Family and Within Family Results

| Outcome | Education | Education | Education |
| --- | --- | --- | --- |
| Sample | Full | Sibling | Sibling |
| Fixed Effects? | None | None | Sibling |
| PGS (std) | 0.908***<br>(0.008) | 0.959***<br>(0.026) | 0.444***<br>(0.048) |
| Female | -0.746***<br>(0.016) | -0.777***<br>(0.055) | -0.860***<br>(0.070) |
| Age Fixed Effects | X | X | X |
| PC Controls (1-20) | X | X | X |
| Observations | 404,409 | 34,209 | 33,044 |
| R-squared | 0.068 | 0.063 | 0.669 |

Notes: Robust standard errors in parentheses, \*\*\* p<0.01, \*\* p<0.05, \* p<0.1

Sibling-difference analysis controls for influences shared by siblings growing up in the same household, including the portion of the PGS associated with genetic nurture (26, 27) that is shared among siblings. However, unshared environmental factors, including parental actions to reinforce or compensate for sibling differences in PGS are retained in the PGS associations. SI Table 2 presents average differences and correlations in sibling phenotype and genotype measures. As expected, sibling correlations in the genotype measures are r~0.5. Correlations in EA and height are ~0.3. On average siblings differ in PGS by ~0.75 SD, 4 years of schooling, and 8 cm in height.

**Table 2.** Within Family Associations between PGS-Education and Educational Attainment Including Sibling Dyad Measures of Relative Position

| Outcome | Education | Education | Education |
| --- | --- | --- | --- |
| Sample | Sibling | High SES Place | Low SES Place |
| Fixed Effects? | Sibling | Sibling | Sibling |
| PGS (std) | 0.389***<br>(0.082) | 0.433***<br>(0.113) | 0.353***<br>(0.119) |
| Female | -0.857***<br>(0.070) | -0.626***<br>(0.096) | -1.083***<br>(0.103) |
| Older Sib of Pair | 0.443***<br>(0.090) | 0.323***<br>(0.122) | 0.576***<br>(0.133) |
| Sib with larger PGS | 0.124<br>(0.076) | 0.183*<br>(0.106) | 0.078<br>(0.108) |
| Larger X PGS | -0.086*<br>(0.050) | -0.179**<br>(0.070) | -0.006<br>(0.073) |
| Age Fixed Effects | X | X | X |
| PC Controls (1-20) | X | X | X |
| Observations | 33,044 | 16,810 | 16,234 |
| R-squared | 0.669 | 0.669 | 0.641 |

Notes: Robust standard errors in parentheses (clustered by family). High SES Place: Families who grew up in places/years with above median predicted educational attainments. Low SES place: Families who grew up in places/years with below median predicted educational attainments. \*\*\* $p < 0.01$ , \*\* $p < 0.05$ , \* $p < 0.1$

In order to conduct an omnibus test of reinforcing vs. compensating processes in reaction to sibling differences in PGS, we add dyadic measures of relative ranking of PGS and age (see Methods) as well as an interaction between the indicator for higher relative ranking and PGS to predict EA.

Table 2 shows that inclusion of the dyadic measures is consequential. Conforming to the birth order literature, the older sibling (even accounting for age indicators (i.e. fixed effects)) attains over 0.5 years of schooling more than the younger sibling of the pair (28–30).^2^ The interaction between the indicator for higher ranked EA-PGS and the EA-PGS score is negative (p-value <0.09), so that the EA-PGS has a smaller association with attainment for siblings who are higher ranked within the sibship (See Figure 1). This result suggests, on average, compensatory processes within families and a concomitant reduction in within-family inequalities in educational attainment.

**Figure 1.**
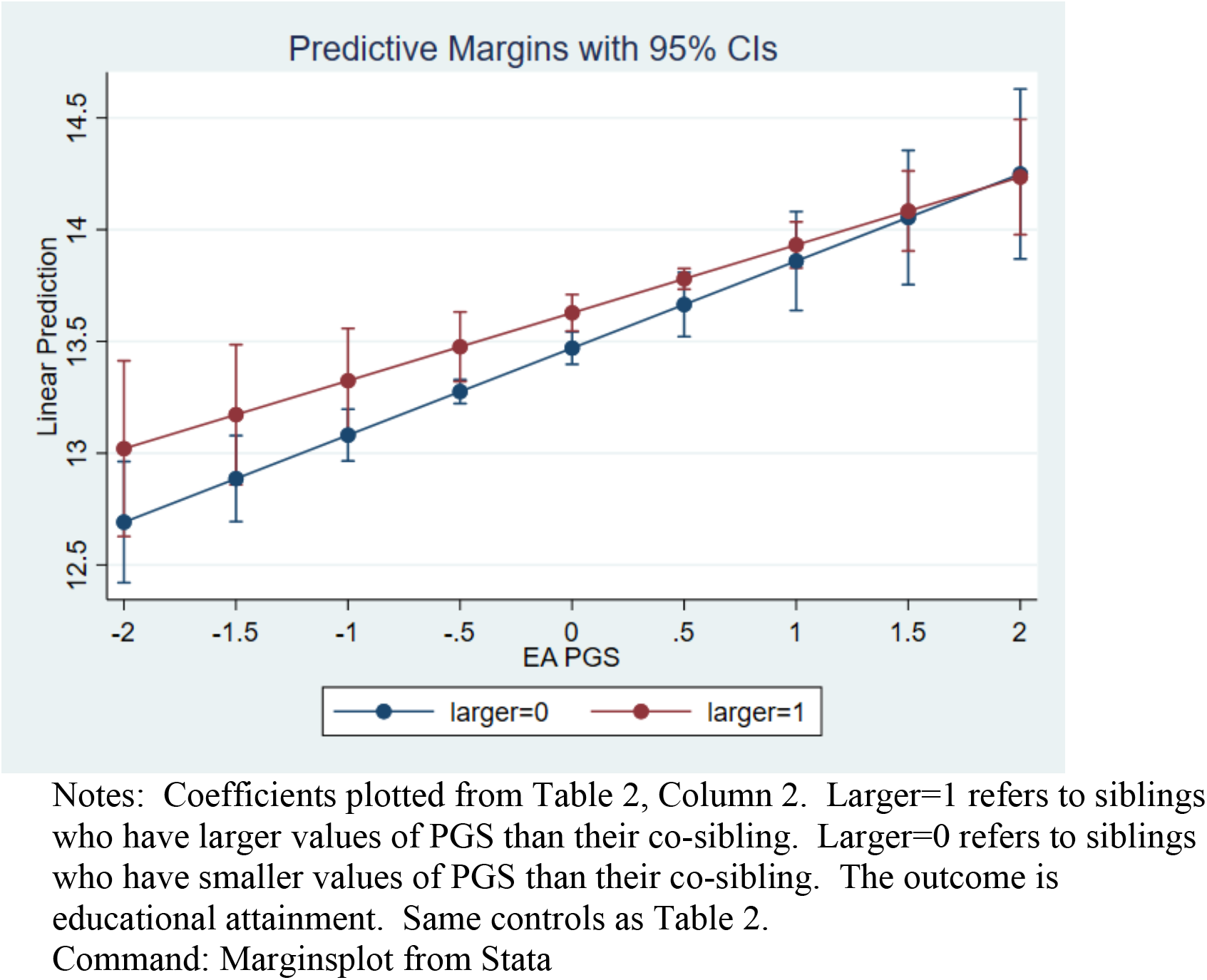
Plot of Associations between EA-PGS and Educational Attainment Stratified by Whether Sibling has larger or smaller PGS than co-Sibling

Table 2 Column 3 and 4 stratify the sibling analysis based on area (place of birth)-estimates of socioeconomic status. Results show that birth order effects are smaller in high SES areas, the gender gap is smaller in high SES areas, and the compensatory effects are much larger in high SES areas (p-value <0.05) (See Figures 2 and 3). These results also show, in general, the importance of families for shaping links between genetic measures and social outcomes.

**Figure 2.**
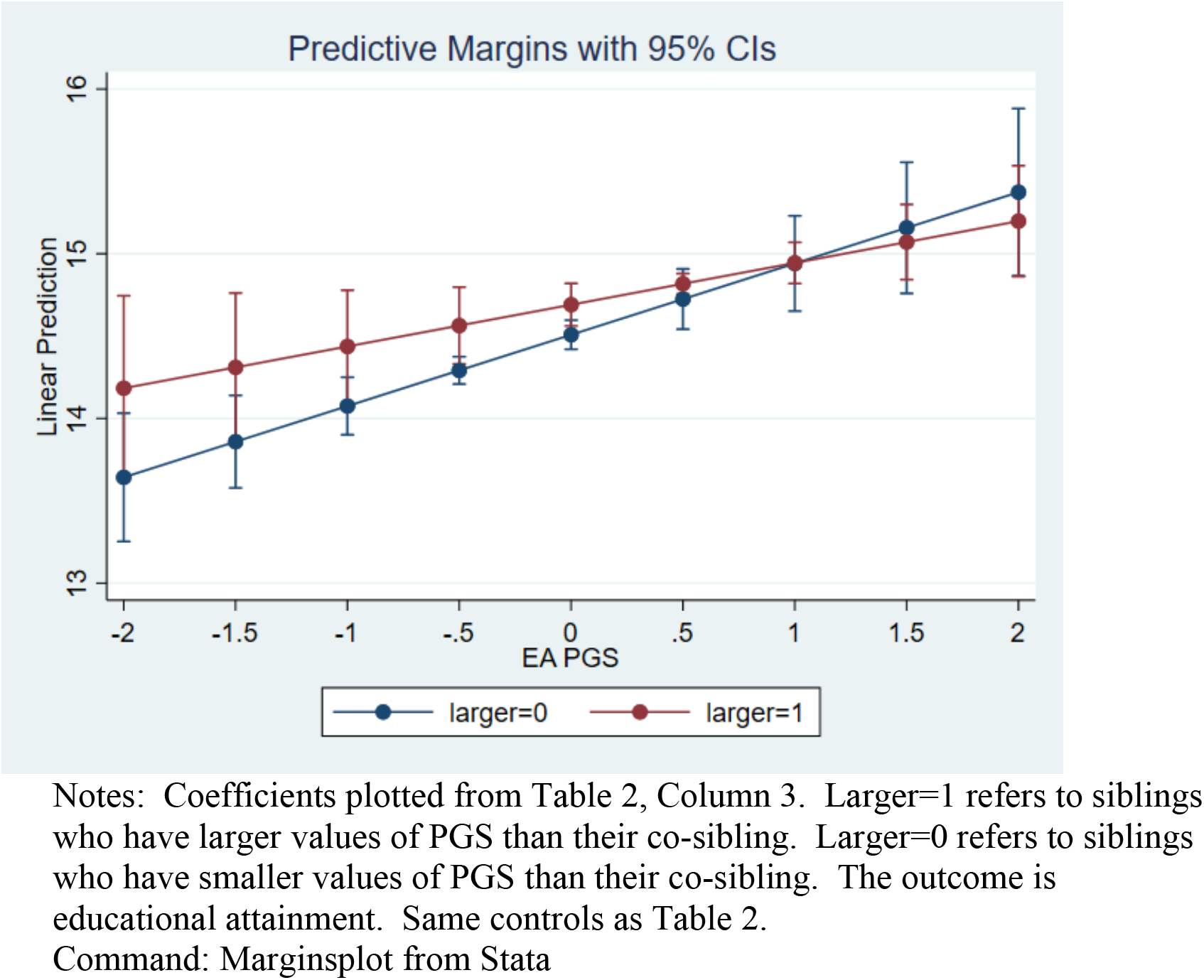
Plot of Associations between EA-PGS and Educational Attainment Stratified by Whether Sibling has larger or smaller PGS than co-Sibling High SES Places

**Figure 1.**
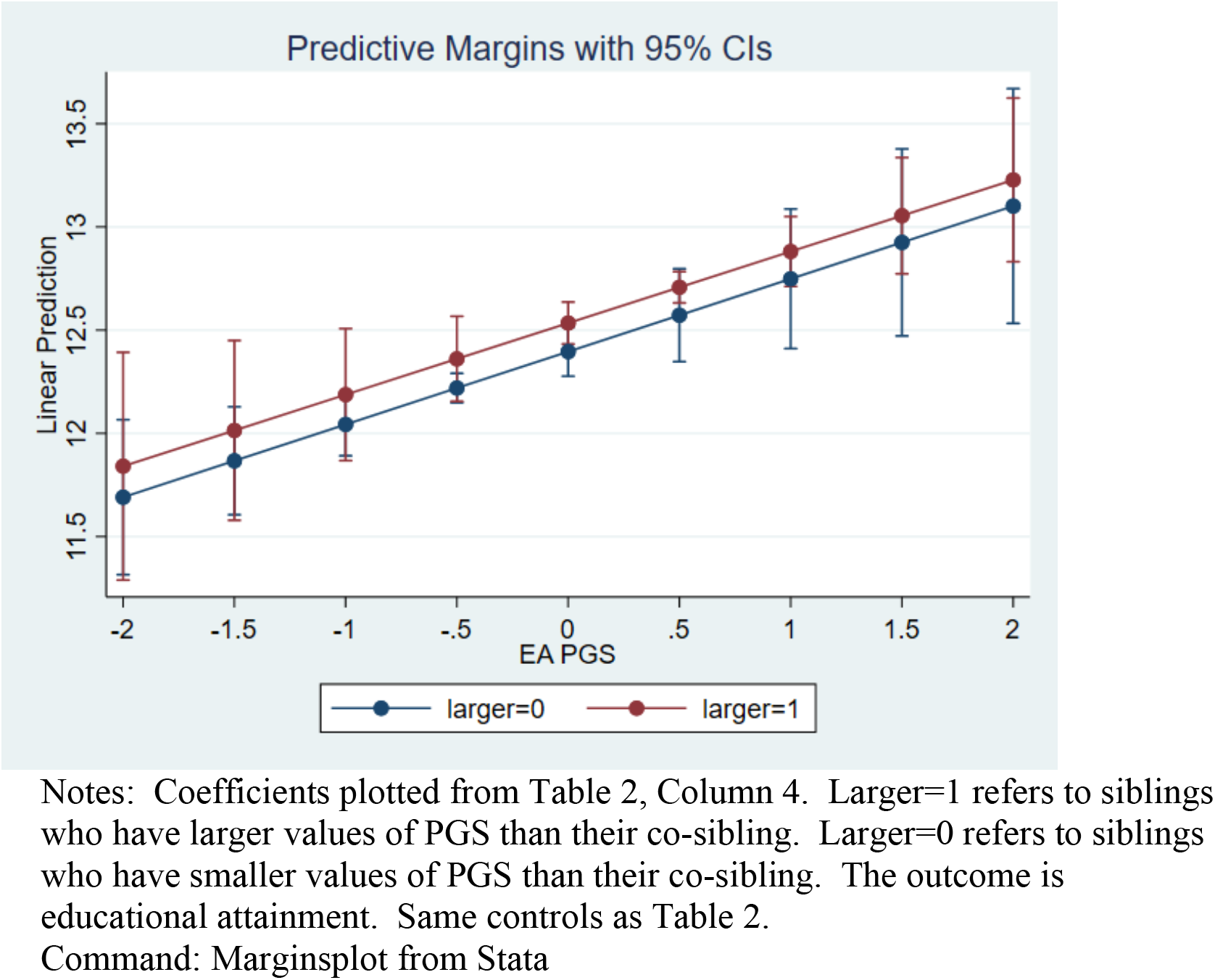
Plot of Associations between EA-PGS and Educational Attainment Stratified by Whether Sibling has larger or smaller PGS than co-Sibling Low SES Places

Table 3 performs a negative test. We hypothesized to find no effect of families and relative rank on genetic penetrance for the case of height. The results suggest no effect and can rule out even modest size effects.

**Table 3:** Falsification Exercise Within Family Associations between PGS-Height and Height

| Outcome | Height | Height | Height | Height |
| --- | --- | --- | --- | --- |
| Sample | Full | Sibling | Sibling | Sibling |
| Fixed Effects? | None | None | Sibling | Sibling |
| Height PGS (std) | 2.516***<br>(0.010) | 2.557***<br>(0.034) | 2.151***<br>(0.050) | 2.154***<br>(0.085) |
| Female | -13.319***<br>(0.019) | -13.295***<br>(0.065) | -13.468***<br>(0.070) | -13.467***<br>(0.070) |
| Older Sib of Pair |  |  |  | 0.019<br>(0.089) |
| Sib with Larger PGS |  |  |  | -0.034<br>(0.075) |
| Larger X PGS |  |  |  | 0.052<br>(0.051) |
| Age Fixed Effects | X | X | X | X |
| PC Controls (1-20) | X | X | X | X |
| Observations | 402,320 | 34,036 | 32,700 | 32,700 |
| R-squared | 0.600 | 0.598 | 0.901 | 0.901 |

Robust standard errors in parentheses (clustered by family), \*\*\* p<0.01, \*\* p<0.05, \* p<0.1

Table 4 performs a second negative test. We form dyads in the data who are related (e.g. cousins) but are not full siblings (kinship <0.18). We show that earlier results of impacts of relative rankings of PGS are absent in these related, non-sibling pairs.

**Table 4:**
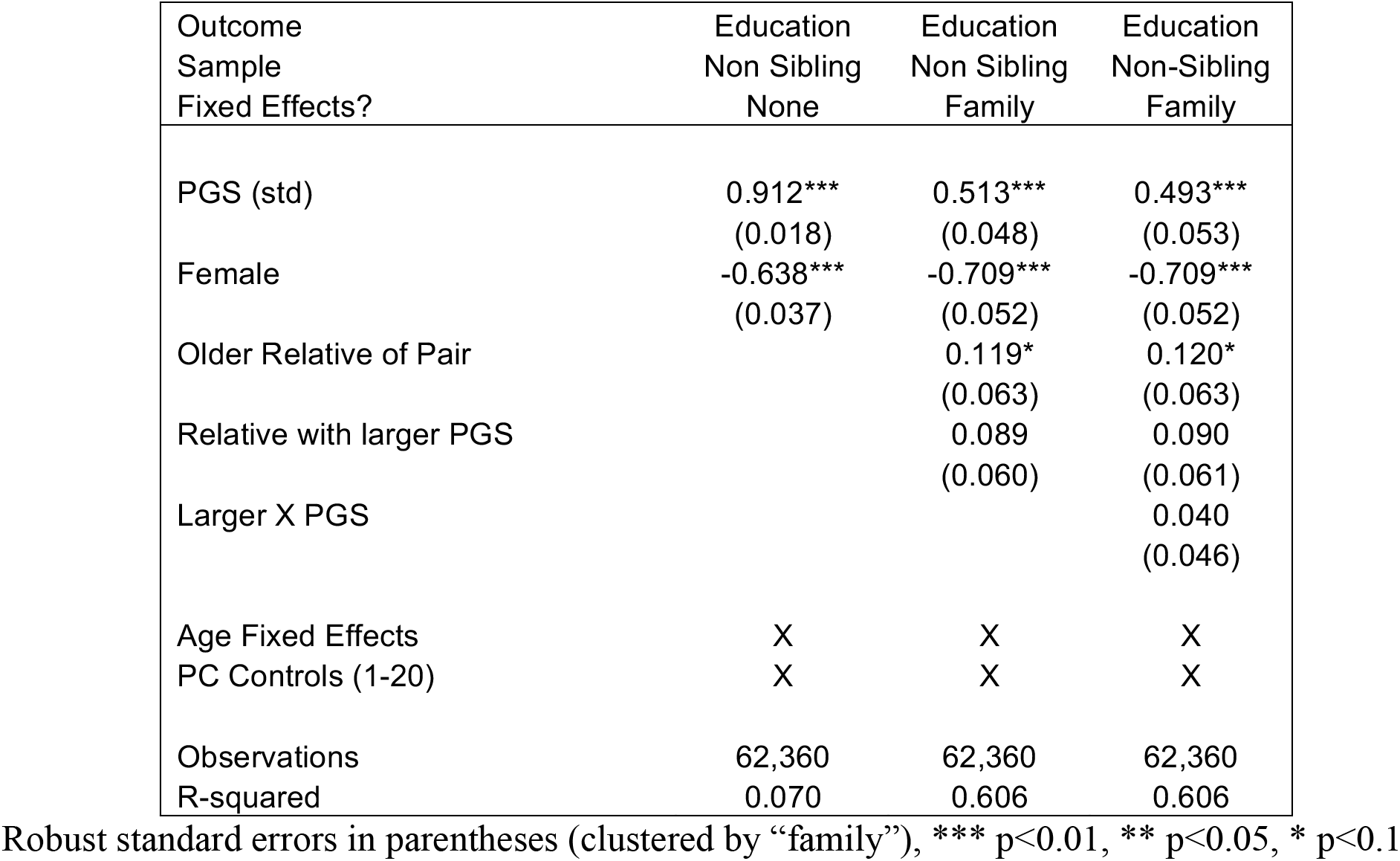
Falsification Exercise Between-Relative Associations of PGS-Education, Educational Attainment, and Relative Rankings of PGS

## Discussion

These results contribute to several literatures in the social and genetic sciences. In the social sciences, the use of PGS to measure relative traits between siblings allows expanded analyses estimating the extent to which parents compensate or reinforce their children’s endowments. Future work could expand this direction by estimating a larger set of domains of PGS as well as focusing on early life measurements of children’s outcomes. A major limitation for this direction of inquiry is the modest sample of siblings in many datasets.

Our results show lower levels of genetic penetrance for educational attainment for children who have higher PGS than their sibling. This phenomenon is larger in places whose residents have higher socioeconomic status than in places with low socioeconomic status. Together, these results are consistent with parental preferences for equality among their children, which may be accomplished through compensating investments.

The findings are also consequential for genetic analyses more broadly. Some recent innovative analysis linking genetics to outcomes has taken seriously the confounding between children’s genetics and family background when explaining children’s later outcomes as adults, seeking to decompose PGS into direct (child) and indirect (parent) effects (26, 31, 32). This work has been motivated, in part, by previous research that used sibling comparisons in PGS to predict education. For example, some early use of educational attainment PGS (33) used sibling pairs to show that the PGS continued to predict education. The implication was then drawn that the PGS contained some causal effects of genetics on important outcomes such as schooling.

Later work has prioritized comparisons of between family estimates with within-family estimates to assure a lack of confounding by family background—using EA2 (34) and EA3(23) has been a bit more mixed, with the within-family analysis often reducing the estimates of genetic penetrance by 50% compared to between-family analysis. However, even with these smaller effects, researchers have relied on this framework to show the likelihood of causal PGS effects on education and related outcomes (25, 35, 36). New work (36) has discussed common issues in genetic analysis that can be overcome with family-based designs, including population stratification, assortative mating, and dynastic effects. One commonality of these issues is that they are presumed to reflect shared effects on all children in a household that can be eliminated through sibling-comparisons or, alternatively, the use of parental genetic data. Indeed, researchers have stated that sibling analysis “rules out purely social transmission as an explanation for the associations between children’s education-linked genetics and their attainment.”(25)

However, while these may be the most common concerns for analysis linking genetics with outcomes, a rich set of theoretical models and empirical results from demography and adjacent social sciences suggest there are yet another set of complications that should be explored before claiming that genetic effects are causal. In particular, families shape outcomes, parents react to children’s abilities and have preferences for their set of children’s outcomes. Other family members (who also often share genetics with the child/children) also contribute. An important outcome of these processes is the possibility of statistical interference between the outcomes of siblings(37), so that siblings designs both eliminate some empirical concerns but raise others.

The current results demonstrate that whatever causal effects exist between PGS and outcomes can be mediated through families, even with a sibling fixed effects design. This suggests caution in interpreting sibling fixed effects models as “causal genetic effects” and reemphasizes that the value of this strategy is to control for shared environments but not produce causal estimates. Indeed, our results suggest that within-sibling analysis of genetics and outcomes continue to reflect family processes rather than pure “genetic effects”. Future work will need to incorporate additional strategies to separate “genetic” and “family” causal processes.

## Methods and Materials

### Data

We used data from the UK Biobank project(38). The participants, aged between 37 and 74 years, were originally recruited between 2006 and 2010. These data are restricted, but one can gain access by following the procedures described in www.ukbiobank.ac.uk/register-apply/. Although siblings are not identified in the survey, respondents’ genetics can be used to measure genetic relatedness among all pairs of respondents. We first use the UKB provided kinship file, listing all pairwise kinships among 100,000 pairs in the sample of nearly 500,000 individuals. We first choose pairs with kinship >0.2, which reflects first degree biological relatives (parents/siblings). We then choose remaining pairs who are <13 years apart in age, leaving ~22,000 sibling dyads. We then chose to keep only one dyad from any family with more one dyad, leaving ~17,600 dyads. We include only respondents of European ancestry in our analysis.

### Polygenic scores

We constructed PGS for two traits for which large genome wide association studies (GWAS) are publicly available and do not contain UKB samples: Height (39) and Educational Attainment(40). We removed single-nucleotide polymorphisms (SNPs) in strong linkage disequilibrium (LD). We LD-clumped the GWAS summary data by PLINK(41), using 1000 Genome Project Phase III European genotype data as reference. We used a LD window size of 1Mb and a pairwise r^2^ threshold of 0.1. We did not apply any p-value thresholding to select SNPs. Final weights were produced by using PRSice-2(42). The PGSs were normalized to have mean zero and SD one and oriented so that each PGS was positively correlated with its corresponding outcome.

### Phenotypes

Educational levels of the UK Biobank participants were measured by mapping each major educational qualification that can be identified from the survey measures to an International Standard Classification of Education (ISCED) category and imputing a years-of-education equivalent for each ISCED category(43). Height is measured standing height.

### Sibling variables

We created three variables indicating the relative status of the members of sibling pairs. First, we created an indicator for the sibling with the higher EA-PGS score. Second, we created an indicator for the sibling with the higher Height-PGS score. Third, we created an indicator for the sibling who is older, due to well known birth order effects on educational attainment.(28–30) We note that self-reported birth order is only available for a subset of UKB respondents.

### Place based socioeconomic status

The UKB does not contain information about childhood background socioeconomic status of the respondents. Therefore we created information based on place of birth and year of birth to predict which places/years of birth had high vs. low (based on median) predicted educational attainment. We split families based on whether neither sibling was born in a place/year with high predicted attainment vs. either sibling was born in a place/year with high predicted attainment. The median predicted schooling was 13.65 years in a regression that had place of birth fixed effects and year of birth fixed effects.

### Sample Characteristics

SI Table 1 presents descriptive statistics for the full sample and the analysis sub-sample of siblings. Educational attainment is slightly higher in the full sample and number of siblings (by definition) is higher in the sibling sample. EA3 is slightly higher on average in the sibling sample but demographic characteristics are quite similar.

### Sample Characteristics for Falsification Exercise

We created a second sample of related individuals who were not full biological siblings in order to develop a negative (falsification) test of the analysis. To construct the sample, we used dyads with kinship measurements below 0.18 and randomly chose one dyad in cases where a respondent was linked with multiple sample members. Sample characteristics are shown in SI Table 3A and Dyadic Characteristics are shown in SI Table 4A.

### Analysis

We tested associations using linear regression models. We clustered standard errors at the family level. We conducted sibling difference analysis using family fixed effects regression (44). We control for sex, age indicators, and 20 genetic principal components. We controlled for age indicators (fixed effects) for the well documented secular increases in schooling over this time period.(45)

We use regression analysis to link PGS to educational attainments in our results. We compare results with and without sibling fixed effects and also with measures reflecting siblings’ relative position in the dyad based on EA3 and age. That is, we regress educational attainment for respondent *i* in family *f* on demographic characteristics (age, sex), a polygenic score and controls for 20 principle components and a family-clustered error term:

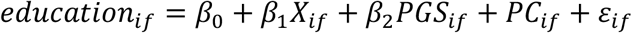

We next add sibling fixed effects to the model.

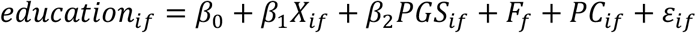

Finally, we enrich the model with dyadic measures of relative position, namely an indicator for whether the respondent has a higher PGS than his/her sibling and the interaction between this indicator and PGS:

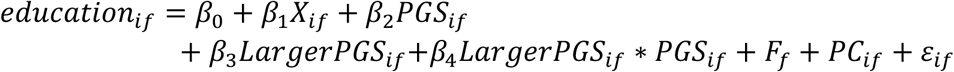

Where the key coefficient is *β*_4_, which reflects differences in genetic penetrance based on relative ranking in the sibship on the PGS. A positive coefficient would suggest that parents reinforce early evidence of a child’s ability (relative to his/her sibling) while a negative coefficient suggests that parents compensate. It is possible that these effects vary by family socioeconomic status.^3^ Therefore, we stratify the model between families who were born in high socioeconomic status areas/years vs. low socioeconomic status areas/years.

## Supplemental Information

### Appendix Tables

**Table 1A.** Comparison of Full UKB Sample and Sibling Analysis Sample

| Variable | Full Sample |  |  |  |  | Sibling Sample |  |  |  |  |
| --- | --- | --- | --- | --- | --- | --- | --- | --- | --- | --- |
|  | Obs | Mean | Std Dev | Min | Max | Obs | Mean | Std Dev | Min | Max |
| Education | 399,442 | 13.8 | 5.1 | 7.0 | 20.0 | 33,815 | 13.6 | 5.1 | 7.0 | 20.0 |
| Female | 403,183 | 0.54 | 0.5 | 0.0 | 1.0 | 34,095 | 0.58 | 0.5 | 0.0 | 1.0 |
| Age | 403,183 | 56.8 | 8.0 | 39.0 | 73.0 | 34,095 | 57.0 | 7.3 | 40.0 | 70.0 |
| EA3 PGS (std) | 403,183 | 0.00 | 1.0 | -4.7 | 5.0 | 34,095 | 0.05 | 1.0 | -3.7 | 4.0 |
| Number of Siblings | 397,344 | 2.0 | 1.8 | 0.0 | 32.0 | 34,019 | 2.8 | 1.9 | 1.0 | 22.0 |
| PC 1 | 403,183 | -12.4 | 1.6 | -18.5 | -6.1 | 34,095 | -12.4 | 1.6 | -18.3 | -6.1 |
| PC 2 | 403,183 | 3.8 | 1.5 | -2.0 | 9.5 | 34,095 | 3.8 | 1.5 | -1.7 | 9.4 |
| PC 3 | 403,183 | -1.6 | 1.6 | -7.7 | 4.4 | 34,095 | -1.6 | 1.6 | -7.4 | 4.3 |
| PC 4 | 403,183 | 1.3 | 2.9 | -10.1 | 12.8 | 34,095 | 1.5 | 2.9 | -9.8 | 12.5 |
| PC 5 | 403,183 | -0.7 | 6.7 | -17.7 | 24.8 | 34,095 | -0.5 | 6.9 | -16.3 | 24.7 |
| PC 6 | 403,183 | -0.4 | 1.6 | -6.7 | 6.0 | 34,095 | -0.4 | 1.6 | -6.6 | 5.9 |
| PC 7 | 403,183 | 0.3 | 1.9 | -17.7 | 11.5 | 34,095 | 0.3 | 1.8 | -13.8 | 7.2 |
| PC 8 | 403,183 | -0.5 | 2.0 | -9.0 | 16.7 | 34,095 | -0.5 | 1.9 | -8.7 | 16.7 |
| PC 9 | 403,183 | 0.0 | 4.6 | -36.6 | 12.6 | 34,095 | 0.3 | 4.4 | -32.4 | 11.4 |
| PC 10 | 403,183 | 0.2 | 2.3 | -11.3 | 11.4 | 34,095 | 0.2 | 2.2 | -9.8 | 9.8 |
| PC 11 | 403,183 | 0.0 | 3.6 | -11.0 | 20.3 | 34,095 | 0.0 | 3.5 | -10.0 | 16.6 |
| PC 12 | 403,183 | 0.2 | 2.3 | -11.3 | 10.9 | 34,095 | 0.2 | 2.3 | -9.2 | 9.0 |
| PC 13 | 403,183 | 0.0 | 1.6 | -7.2 | 7.2 | 34,095 | 0.0 | 1.6 | -7.2 | 7.0 |
| PC 14 | 403,183 | 0.2 | 3.3 | -14.1 | 18.2 | 34,095 | 0.2 | 3.2 | -11.5 | 17.3 |
| PC 15 | 403,183 | 0.0 | 1.9 | -13.8 | 9.6 | 34,095 | 0.0 | 1.8 | -10.8 | 9.0 |
| PC 16 | 403,183 | 0.0 | 3.1 | -27.8 | 20.5 | 34,095 | 0.1 | 3.0 | -22.7 | 16.7 |
| PC 17 | 403,183 | 0.0 | 2.0 | -9.9 | 8.7 | 34,095 | 0.0 | 1.9 | -9.1 | 7.9 |
| PC 18 | 403,183 | 0.0 | 2.9 | -16.6 | 21.2 | 34,095 | 0.0 | 2.8 | -12.2 | 15.9 |
| PC 19 | 403,183 | 0.0 | 2.9 | -11.7 | 14.6 | 34,095 | 0.0 | 2.7 | -9.7 | 11.6 |
| PC 20 | 403,183 | 0.0 | 2.9 | -12.6 | 14.9 | 34,095 | 0.1 | 2.7 | -10.9 | 12.9 |

**Table 2A.** Dyadic (Siblings) Measures Differences and Correlations

| Variable | Obs | Differences |  |  |  | Correlations |
| --- | --- | --- | --- | --- | --- | --- |
|  |  | Mean | Std Dev | Min | Max |  |
| EA PGS (Std) | 16,792 | 0.77 | 0.59 | 0.0 | 4.3 | 0.54 |
| Height PGS (Std) | 16,792 | 0.75 | 0.57 | 0.0 | 4.1 | 0.57 |
| PC1 | 16,792 | 1.21 | 0.93 | 0.0 | 6.0 | 0.54 |
| Education | 16,522 | 4.14 | 4.24 | 0.0 | 13.0 | 0.32 |
| Height | 16,734 | 8.68 | 6.62 | 0.0 | 60.0 | 0.29 |
| Age | 16,792 | 4.25 | 2.75 | 0.0 | 12.0 | 0.76 |

**Table 3A.** Descriptive Statistics for Relatives (Non-Full-Siblings) Sample

| Variable | Obs | Mean | Std Dev | Min | Max |
| --- | --- | --- | --- | --- | --- |
| Education | 62,360 | 13.19 | 5.12 | 7.00 | 20.00 |
| Female | 62,360 | 0.54 | 0.50 | 0.00 | 1.00 |
| Age | 62,360 | 56.96 | 7.94 | 39.00 | 70.00 |
| EA3 PGS (std) | 62,360 | -0.02 | 1.00 | -4.57 | 4.08 |
| Height PGS (std) | 62,360 | -0.02 | 1.00 | -4.40 | 4.19 |
| Height (cm) | 62,229 | 168.37 | 9.22 | 100.00 | 204.00 |
| Larger EA3 PGS than Relative | 62,360 | 0.50 | 0.50 | 0.00 | 1.00 |
| Older Than Relative | 62,360 | 0.50 | 0.50 | 0.00 | 1.00 |
| Number of Siblings | 61,312 | 2.55 | 1.86 | 1.00 | 23.00 |
| PC 1 | 62,360 | -12.40 | 1.59 | -18.43 | -6.10 |
| PC 2 | 62,360 | 3.79 | 1.50 | -1.91 | 9.50 |
| PC 3 | 62,360 | -1.62 | 1.56 | -7.39 | 4.30 |
| PC 4 | 62,360 | 1.53 | 2.90 | -10.14 | 12.46 |
| PC 5 | 62,360 | -0.37 | 6.78 | -16.02 | 24.79 |
| PC 6 | 62,360 | -0.38 | 1.61 | -6.72 | 5.94 |
| PC 7 | 62,360 | 0.27 | 1.82 | -17.67 | 8.27 |
| PC 8 | 62,360 | -0.46 | 1.93 | -8.44 | 16.07 |
| PC 9 | 62,360 | 0.18 | 4.37 | -33.88 | 11.36 |
| PC 10 | 62,360 | 0.19 | 2.19 | -9.10 | 10.72 |
| PC 11 | 62,360 | 0.08 | 3.43 | -10.62 | 17.37 |
| PC 12 | 62,360 | 0.14 | 2.28 | -10.88 | 9.56 |
| PC 13 | 62,360 | 0.03 | 1.59 | -6.88 | 6.78 |
| PC 14 | 62,360 | 0.21 | 3.15 | -11.91 | 18.16 |
| PC 15 | 62,360 | 0.00 | 1.82 | -12.26 | 8.53 |
| PC 16 | 62,360 | -0.02 | 2.97 | -22.79 | 18.16 |
| PC 17 | 62,360 | 0.00 | 1.87 | -8.24 | 8.43 |
| PC 18 | 62,360 | 0.10 | 2.80 | -12.22 | 17.61 |
| PC 19 | 62,360 | -0.05 | 2.71 | -11.39 | 11.43 |
| PC 20 | 62,360 | 0.13 | 2.70 | -12.45 | 13.17 |

**Table 4A.**
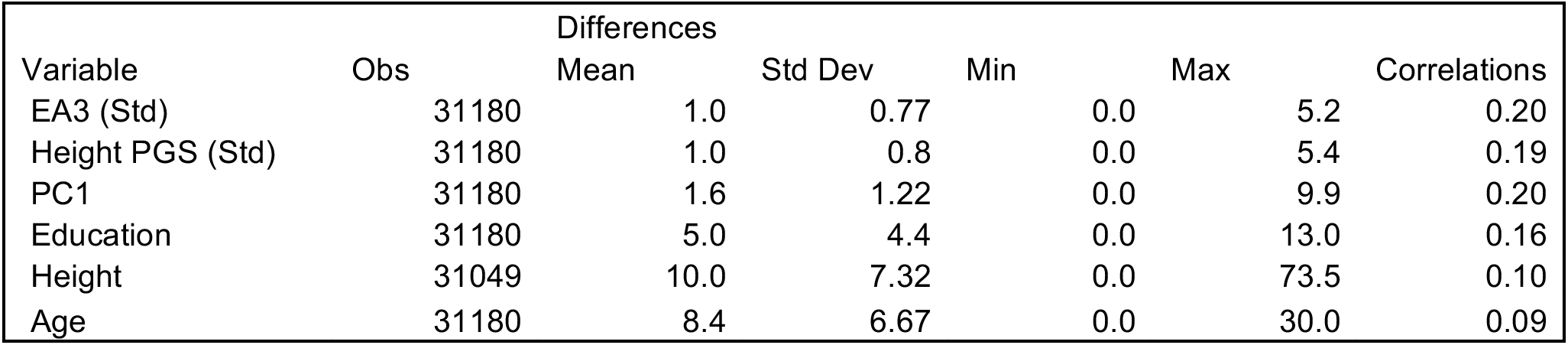
Dyadic (Relatives, Non Full Siblings) Measures Differences and Correlations

| Variable | Obs | Differences |  | Min | Max | Correlations |
| --- | --- | --- | --- | --- | --- | --- |
|  |  | Mean | Std Dev |  |  |  |
| EA3 (Std) | 31180 | 1.0 | 0.77 | 0.0 | 5.2 | 0.20 |
| Height PGS (Std) | 31180 | 1.0 | 0.8 | 0.0 | 5.4 | 0.19 |
| PC1 | 31180 | 1.6 | 1.22 | 0.0 | 9.9 | 0.20 |
| Education | 31180 | 5.0 | 4.4 | 0.0 | 13.0 | 0.16 |
| Height | 31049 | 10.0 | 7.32 | 0.0 | 73.5 | 0.10 |
| Age | 31180 | 8.4 | 6.67 | 0.0 | 30.0 | 0.09 |

## Footnotes

2 We would have preferred to control for birth order, but the UKB only asked birth order information for a subset of the sample.

3 Unfortunately, the UKB is extremely limited in measurements of childhood experiences and has nearly no information about parents’ characteristics of the respondents (e.g. parental education, family socioeconomic status during childhood, etc).

